# Decreasing ELK3 expression improves Bone Morphogenetic Protein Receptor 2 signaling and pulmonary vascular cell function in PAH

**DOI:** 10.1101/2023.01.14.524023

**Authors:** Md Khadem Ali, Lan Zhao, Vinicio de Jesus Perez, Mark R. Nicolls, Edda F. Spiekerkoetter

## Abstract

ELK3 is upregulated in blood and pulmonary vascular cells of PAH patients and may play a significant role in PAH potentially through modulating BMPR2 signaling.

## Introduction

Pulmonary arterial hypertension (PAH) is a rare but complex, severe, life-threatening condition of the small pulmonary arteries. PAH is characterized by pulmonary arterial remodeling and increased pulmonary vascular resistance; the consequent elevated pulmonary arterial pressures causes right ventricular afterload and, ultimately, failure. The disease remains incurable despite intense efforts to identify new therapies to treat PAH patients. In order to identify new therapies, it is crucial to understand the cause and exact molecular mechanisms of the disease. Although the exact cause of PAH is unclear, many genetic, epigenetic, and environmental factors have been shown to contribute to the development and progression of the disease. For example, heterozygous loss of function mutation in bone morphogenic protein receptor 2 (BMPR2) occurs in 53-86% of familial PAH patients and 14-35% in sporadic idiopathic PAH patients(1). However, the disease penetrance rate of the mutation carriers is low, indicating that other unidentified factors may contribute to the disease development in addition to the gene mutation. Significantly, BMPR2 signaling is thought to be impaired in PAH patients regardless of the etiology of PAH, making the signaling a master switch in the disease. Previously, we and others have shown that targeting BMPR2 signaling with repurposed drugs (FK506, Enzastaurin) or rebalancing the pathway with BMP9 and Sotatercept improved PAH in animal models and pilot studies in patients (1).

To identify BMPR2 signaling modifier genes, our team previously performed an siRNA-mediated high throughput screening (HTS) of ∼22,000 genes in a BRE-ID1 incorporated mouse myoblastoma reporter cell line, which yielded two important novel BMPR2 modifier genes (FHIT, LCK) in PAH (2). In the HTS data set, we also found that E26 transformation-specific transcription factor (ELK3) knockdown decreased Id1 levels. Surprisingly, our validation experiments in human pulmonary arterial endothelial cells (PAECs) and pulmonary arterial smooth muscle cells (PASMCs) showed that ELK3 modulated BMPR2 signaling in the opposite direction compared to the HTS experiments using the mouse myoblastoma cell line. ELK3 knockdown increased BMPR2 signaling. ELK3, also known as NET/SAP-2/ERP, is a transcription factor which can form a ternary complex with serum response factor and DNA. While ELK3 generally acts as a transcriptional repressor, it can also work as a transcriptional activator when phosphorylated by the Ras/mitogen-activated protein kinase signaling pathway. ELK3 regulates various biological processes in health and disease, such as proliferation, apoptosis, migration, and angiogenesis. Elevated expression of ELK3 plays a significant role in accelerating the progression and metastasis of different cancers, such as prostate, breast, bladder, gastric, and liver cancer. A significant upregulation of ELK3 expression was also observed in rat carotid arteries following balloon-injury and in human plaques (3). ELK3 was also found to attenuate angiogenesis in VEGF-induced angiogenesis assays *in vitro* and *in vivo* (4). While the role of ELK3 has been studied in different cancers and cardiovascular diseases, its role in PAH is unknown.

## Results and Discussion

We first checked whether ELK3 expression is dysregulated in the whole blood of PAH patients using a large RNA sequencing (RNAseq) data set comprising 359 patients with idiopathic, heritable, and drug-induced PAH as well as 72 age- and sex-matched healthy (5). We found a significant upregulation of ELK3 in the blood of PAH patients compared to healthy controls (**Figure 1A)**. We next tried to determine whether the upregulation of ELK3 in peripheral blood is mirrored by an ELK3 upregulation in the PAH pulmonary vasculature. Thus, we re-analyzed publicly available RNAseq data sets generated from two critical pulmonary vascular cell types in PAH, PAECs and PASMCs of PAH patients and healthy controls. We found a significant increase in ELK3 expression in the PASMCs of 4 idiopathic (I)PAH patients compared to 4 healthy control PASMCs (**Figure 1B**, GSE144274). We also measured the expression of ELK3 in PASMCs of a different cohort of 4 healthy controls and 3 PAH patients by quantitative reverse transcription PCR (qRT-PCR) but did not observe a significant change in ELK3 expression between the two groups (**Figure 1C**). Possible confounding factors are low ELK3 mRNA expression, low sample size and different PAH etiologies, which warrants further validation in a larger cohort. Previously, several studies showed increased expression of ELK3 in PAECs of patients. In a single-cell RNAseq analysis study of lung tissues from 6 healthy controls and 4 IPAH patients, ELK3 expression was shown to be up-regulated in endothelial cells of IPAH patients compared to healthy controls(6). As another example, Reyes-Palomares and colleagues identified that the transcriptional activity of ELK3 was more active in PAH patients than healthy controls (7). We therefore analyzed ELK3 expression in PAECs of PAH patients using a publicly available RNAseq data set comprising 9 healthy controls and 8 PAH patients (GSE0126262). We did not observe a significant change in ELK3 expression in PAECs of PAH patients in this RNAseq dataset (**Figure 1D**). The increased detection of ELK3 in blood of PAH patients could not be attributed to PASCMs and PAECs in our relatively small sample size, given that these cells could be the source of ELK3 in blood but that was not demonstrated in this study.

**Figure 1.**
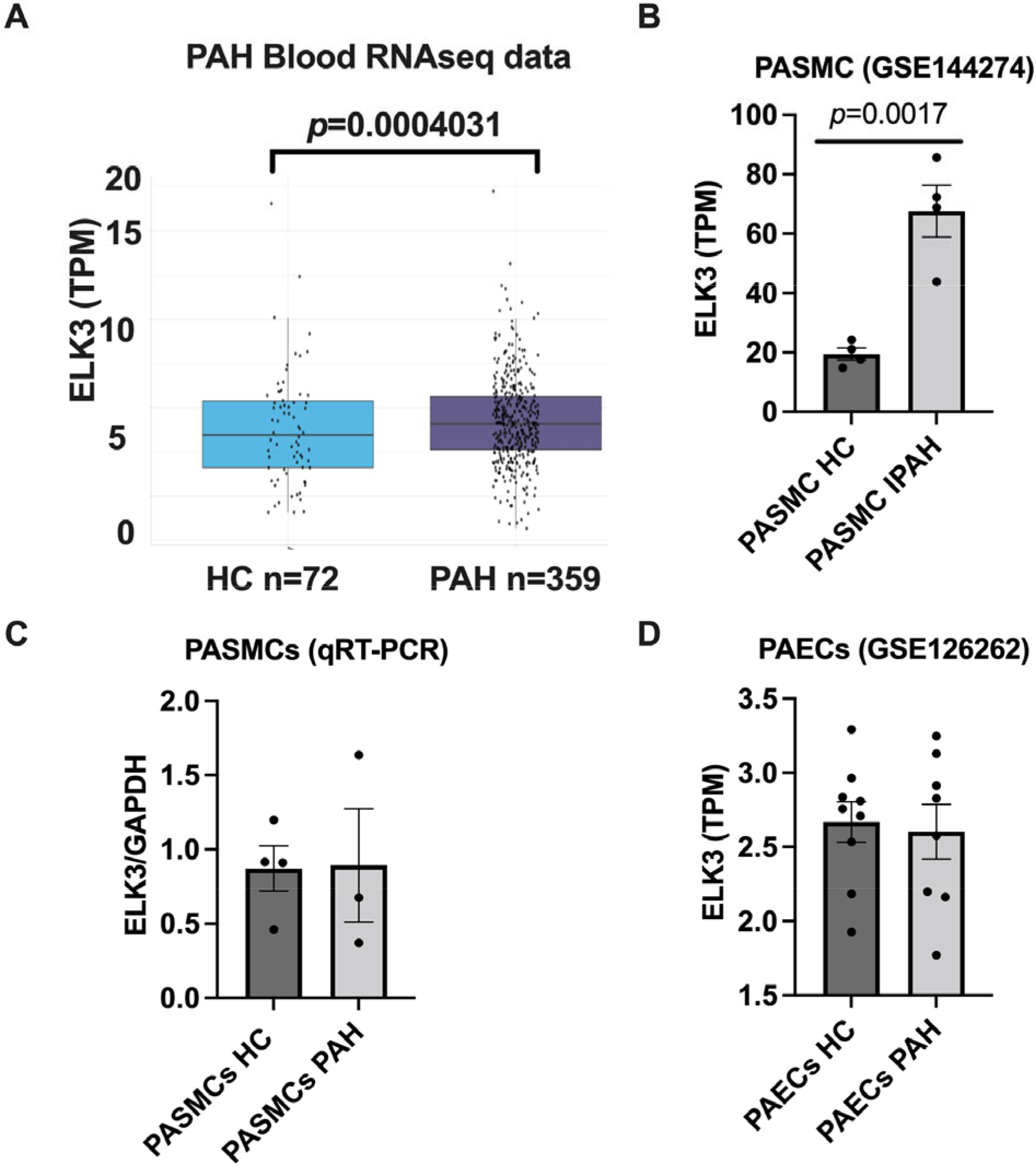
ELK3 is upregulated in the blood and PASMCs of PAH patients. A) RNAseq analysis of the whole blood collected from the 72 healthy controls (HC) and 359 patients with IPAH, APH or HPAH showed a significant increase in ELK3 expression in PAH compared to HC. For detail subject characteristics and hemodynamic data, please see (5). B) ELK3 expression was found to be upregulated in PASMCs of IPAH patients in a publicly available RNAseq data seta comprising 4 HC and 4 IPAH patients (GSE144274). C) ELK3 expression was not altered in PASMCs of a small cohort of PAH patients by qRT-PCR. D) ELK3 expression was not changed in PAECs of a small cohort of PAH patients (GSE126262). Wilcoxon rank sum test with continuity correction was performed to compare ELK3 expression in the whole blood RNAseq data. Student t-test was used to compare ELK3 expression in the PASMCs of PAH RNAseq data. **P<0.01. TPM, transcripts per million.

ELK3 is predicted to be connected with the inhibitor of DNA (ID) signaling pathway (wikipathways). Furthermore, homozygous deficiency in ELK3 was shown to up-regulate expression of SERPINE1 (Serpin family E member 1), also called plasminogen activator inhibitor 1 (PAI-1) in prostate cancer cells(8). PAI-1 is a known downstream target of the BMP signaling pathway and is strongly linked to PAH(9). A recent integrated bioinformatic analysis revealed SERPINE1 (PAI-1) as one of the most significant markers in PAH(10). We therefore hypothesized that the observed increase in ELK3 might downregulate BMPR2 signaling and thereby be involved in PAH pathogenesis. We further hypothesized that decreasing ELK3 might do the opposite – increase BMPR2 signaling – and thereby might be beneficial in PAH. We conducted experiments in PAECs subjected to ELK3 or non-target siRNA with and without BMP9 stimulation. We demonstrated that ELK3 silencing further increased BMP9-induced phospho-SMAD1/5/9 levels measured by western blot (**Figures 2A-B**). ELK3 protein concentration is very low in PASMCs, as seen in **Figure 2C**, and therefore we used qRT-PCR to assess the effect of ELK3 knockdown on BMPR2 signaling. We found a significant increase in BMPR2 as well as ID1 expression, a downstream target of the BMPR2 signaling, following the knockdown of ELK3 with siRNA in PASMCs, while BMPR2 knockdown did not change the ELK3 expression (**Figures 2D-G**), suggesting ELK3 to be upstream of BMPR2. We also observed that inhibition of ELK3 decreased PASMCs proliferation **(Figure 2H)**, which is in line with the proposed model that ELK3 inhibition improves BMPR2 signaling and function in PAEC and PASMC **(Figure 2I)**.

**Figure 2.**
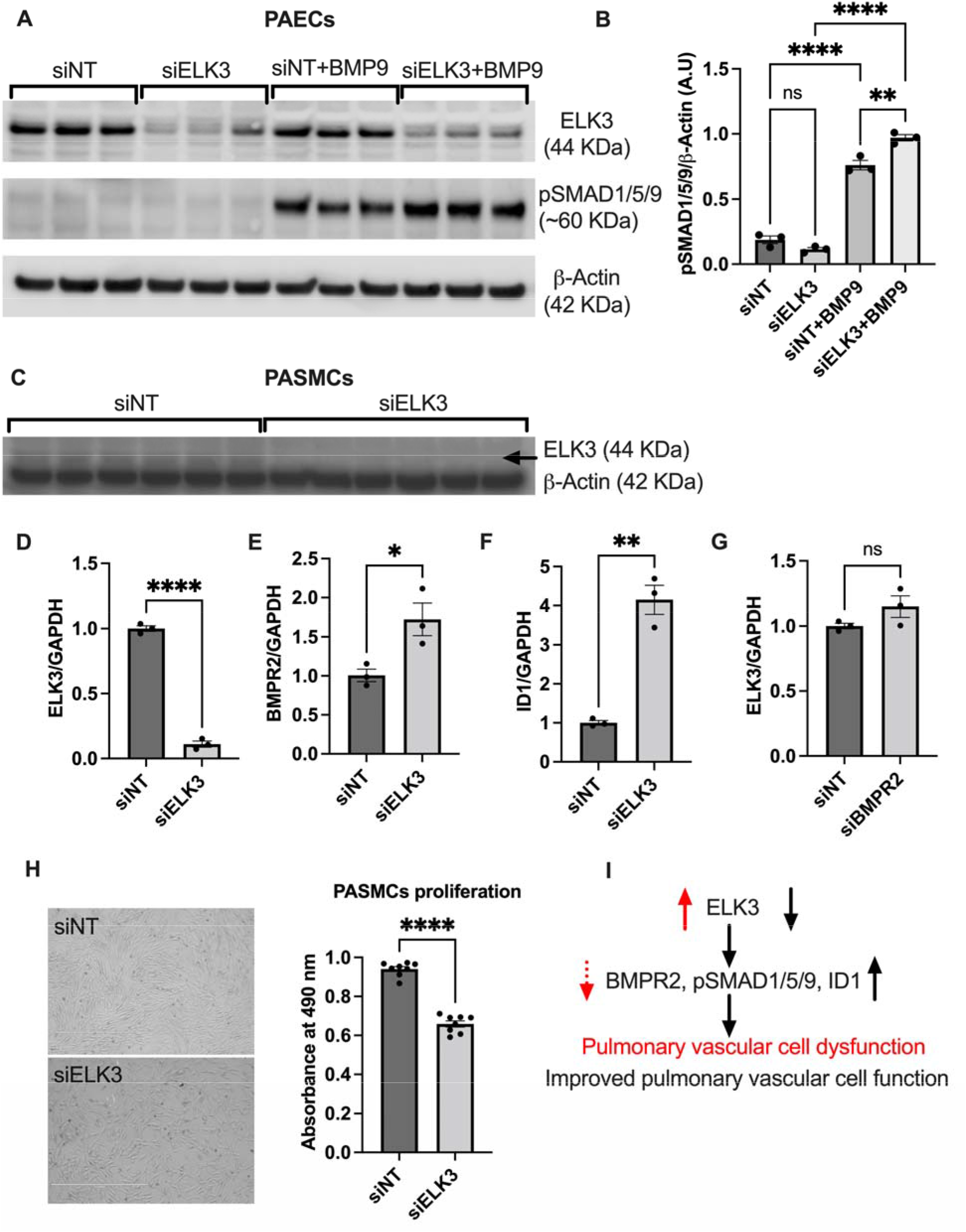
ELK3 is involved in regulating BMPR2 signaling and PASMCs proliferation. A-B) Inhibition of ELK3 with siRNA increased BMP9-induced pSMAD1/5/9 in PAECs, as measured by western blot, 48h knock down, 2h BMP9 stimulation. C) Western blot verification of ELK3 knockdown with siRNA in PASMCs. D-G) siRNA-mediated knockdown of ELK3 increased ID1 levels in PASMCs. BMPR2, ID1 and ELK3 levels were measured by qRT-PCR following 72 hours of ELK3 or BMPR2 knockdown with 60nM siRNA and 3ul RNAimax. Data are represented as mean +/-standard error mean (n=6/group). Student t test, **P<0.01. H) Silencing of ELK3 decreases hPASMCs proliferation as measured by MTT assay. Student t test, ****P<0.0001. I) Proposed model for how ELK3 regulates BMPR2 signaling and PAH.

Previous studies suggested that ELK3 is downregulated during hypoxia, releasing repression of several genes and leading to increased expression of Egr1 and VEGF, as well as PHD2, PHD3, and Siah2 destabilizing HIF1α (6). Therefore, its enhanced expression in PAH is surprising and might be expected to inhibit angiogenesis. Further studies need to explore the exact role and molecular mechanisms of ELK3 in PAH.

This study has several limitations. *First*, we were not able to validate the RNAseq expression of ELK3 by qRT-PCR. *Second*, while ELK3 inhibition increases BMPR2 signaling, it is critical to explore whether overexpression of ELK3 does the opposite, that is, inhibits BMPR2 expression and signaling and induces PAH. *Third*, the molecular mechanisms of how ELK3 regulates BMPR2 signaling, and PAH still need to be clarified.

## Conclusion

In summary, through preliminary experimental and clinical sample analysis, we uncovered ELK3 as a clinically meaningful BMPR2 signaling modulator that influences pulmonary vascular cell function. Further studies are required to fully elucidate the role and molecular mechanisms of ELK3 expression in the pathogenesis of PAH.

## Acknowledgment

The authors gratefully acknowledge the participation of patients recruited to the UK PAH Cohort Study consortium. The authors also thankful to Dr. Christopher J. Rhodes and Professor Martin R. Wilkins at the National Heart and Lung Institute, Hammersmith Campus, Imperial College London, London, UK for their critical comments and statistical analysis of the ELK3 expression in the whole blood of PAH patients and healthy controls by RNAseq.

## Author contribution

MKA and ES conceptualised the study design. MKA performed the experiments and data analysis. All authors contributed to data collection, data interpretation, writing, and editing the manuscript. ES: fund acquisition and supervision.

